# MOTOR MODULES ARE LARGELY UNAFFECTED BY PATHOLOGICAL WALKING BIOMECHANICS: A SIMULATION STUDY

**DOI:** 10.1101/2024.04.08.588563

**Authors:** Mohammad Rahimi Goloujeh, Jessica L. Allen

**Author notes:** Correspondence should be addressed to: Jessica L. Allen, Mechanical and Aerospace Engineering Department University of Florida, PO Box 116250, Gainesville, FL, 32611.

## Abstract

**Background:** Motor module (a.k.a. muscle synergy) analysis has frequently been used to provide insight into changes in muscle coordination associated with declines in walking performance, to evaluate the effect of different rehabilitation intervention, and more recently, to control exoskeletons and prosthetic devices. However, it remains unclear whether changes in muscle coordination revealed via motor module analysis stem from pathological walking biomechanics or pathological neural control. This distinction has important implications for the use of motor module analysis for rehabilitation interventions and device design. Thus, this study aims to elucidate the extent to which motor modules emerge from pathological walking biomechanics.

**Methods:** We conducted a series of computer simulations using OpenSim Moco to simulate abnormal biomechanics by manipulating speed, asymmetry, and step width in a three-dimensional musculoskeletal model. We extracted motor modules using nonnegative matrix factorization from the muscle activation from each simulation. We then examined how alterations in walking biomechanics influenced the number and structure of extracted motor modules and compared the findings to previous experimental studies.

**Results:** The motor modules identified from our simulations were similar to those identified from previously published experiments of non-pathological walking. Moreover, our findings indicate that the same motor modules can be used to generate a range of pathological-like waking biomechanics by modulating their recruit timing over the gait cycle. These results contrast with experimental studies in which pathological-like walking biomechanics are accompanied by a reduction in motor module number and alterations in their structure.

**Conclusions:** This study highlights that pathological walking biomechanics do not necessarily require pathological motor modules. In other words, changes in number and structure of motor modules can be a valuable indicator of alterations in neuromuscular control and may therefore be useful for guiding rehabilitation interventions and controlling exoskeletons and prosthetic devices in individuals with pathological walking function.

## 1. Background

Motor module (a.k.a. muscle synergy) analysis has emerged over the last few decades as a useful method to characterize the complex coordination of the numerous muscles involved in movements such as walking. Motor modules reflect coordinated patterns of muscle activity that can be flexibly combined to meet the goals of different movement behaviors [1]. It is hypothesized that motor modules reflect an underlying nervous system strategy to overcome the complexity of controlling movement by grouping muscles into functional units. As such, many researchers are using motor modules to evaluate the effect of different rehabilitation interventions on neuromuscular control [2–12] and even to control exoskeletons and prosthetic devices [13–21]. However, since motor modules are identified from experimentally-recorded electromyography (EMG) using numerical decomposition techniques such as principal component analysis or non-negative matrix factorization [1,22,23], there is ongoing debate regarding whether they truly represent an underlying neural strategy or simply emerge from the biomechanics of the recorded movement. Incorporating EMG-derived motor modules into rehabilitation interventions and/or into controllers for exoskeletons and prosthetics introduces additional complexity compared to recording only movement biomechanics. If motor modules are simply emergent from movement biomechanics, then there may be no need to incorporate such complexity into these settings. Therefore, it is critical to understand to what extent EMG-derived motor modules reflect an underlying neural strategy to support their use in identifying neuromuscular deficits limiting walking function, guiding rehabilitation efforts, and facilitating control of exoskeleton and prosthetic devices.

Converging evidence suggests that recruiting a reduced number of motor modules to control walking contributes to pathological walking function. Several studies have identified that between four to six motor modules are needed to describe muscle activity in unimpaired walking [24–26]. Comparatively fewer motor modules are required to describe muscle activity in people with neurological deficits (e.g., stroke [7,24,27,28] cerebral palsy [29]and Parkinson’s disease [30–32] and musculoskeletal conditions such as osteoarthritis [33]). Given that each module in unimpaired walking is organized around producing biomechanical functions such as leg swing control and forward propulsion [34,35], it is not surprising that individuals with a reduced number of motor modules typically walk with pathological walking function. For example, reduced motor module number in stroke survivors is associated with slower walking speeds and more asymmetrical steps [24,27] and an inability to change speed, cadence, step length, and step height [28]. Moreover, increases in motor module number that occur with rehabilitation are associated with improved walking function [10,36]. While the prevailing interpretation of these studies is that a decrease in motor module number causes pathological function, an alternative explanation is that the observed reduction in motor modules may be a consequence of the altered walking biomechanics rather than the cause.

Musculoskeletal modeling and simulation offer a means to disentangle the effects of neural control and biomechanics on motor modules. For example, Falisse et al., demonstrated a reduction in motor module number alone could not produce the crouch gait biomechanics often observed in cerebral palsy [37].

Moreover, Mehrabi et. al., found that normal walking biomechanics could not be achieved with reduced motor module number [38]. Taken together, these studies suggest that although biomechanics may have some influence on motor modules, there remains room for them to have a neural basis. However, a limitation of these studies is their focus on a single gait type, preventing insights into the extent of biomechanics versus neural control on motor module structure across diverse pathological walking biomechanics often observed in motor-impaired populations.

The purpose of this study was to explore the extent to which motor modules are emergent from pathological walking biomechanics. Or in other words, does pathological walking function *require* a reduction in motor module number. Inspired by the work of [26] in which motor modules were extracted from individual muscle-driven simulations of unimpaired walking identical to how motor modules are identified experimentally from EMG, we compared motor modules extracted from over 25 muscle-driven simulations of pathological gait biomechanics. Specifically, we investigated the effect of varying speeds, step length asymmetry, and step widths representative of those observed in pathological populations such as stroke survivors. We hypothesized that motor modules cannot be explained by biomechanics alone and thus reflect to some extent an underlying neural control strategy. Based on this hypothesis, we predicted that the number and structure of motor modules would not differ across simulations with different pathological walking biomechanics.

## 2. Methods

A series of 27 different simulations were designed to model different walking biomechanics commonly encountered in post-stroke gait, including three speeds (0.8, 1.1, and 1.45 m/s), step length asymmetry levels (0, 15, and 30%), and step widths (0.1, 0.2, and 0.3 m). For each simulation, optimal muscle recruitment in a 3D musculoskeletal model over a single gait cycle was identified using OpenSim MoCo [39]. Motor modules were then extracted from each of the 27 different optimal solutions from each leg for the (a) full set of 43 muscles per leg and (b) a reduced set of 8 muscles per leg similar to those in which EMG is typically collected experimentally. Finally, the number and structure of motor modules across different walking behaviors were compared to address our study hypotheses.

### 2.1. Musculoskeletal Model

The musculoskeletal model was a modified and armless version of the 3D OpenSim model originally described in Rajagopal et al. (2016)[40]. The modified model consisted of 14 rigid body segments representing the torso, pelvis, and both legs (femur, tibia, patella, calcaneus, talus, and toes) with a total mass of 73.5 kg and height of 1.75 m. The model had 23 degrees-of-freedom (DoF), with a 6 DoF pelvis as the root segment. The trunk and hip were both modeled with three rotational DoF. The knee was replaced by a pin joint for a single DoF. The ankle, subtalar, and toe joints were also modeled as single DoF joints. The model was controlled by 94 Hill-type musculoskeletal units using the DeGrooteFregly2016Muscle model (43 per leg and 4 on each side of the torso; Table 1). Three torque actuators were included around the torso joint to provide additional stability not provided by the trunk muscles. Passive torques representing forces applied by ligaments, passive tissues, and other joint structures were also added to the model via torsional spring-dampers according to the equations in [41,42]. Ground contact was modeled using six Hunt-Crossley contact spheres on each foot: one of radius 3.5 cm positioned at the bottom of the heel, three of radius 1.5 cm at anterior portion of the calcaneus at toe joint, and two with the radius of 1.5 cm on toes). The stiffness and dissipation coefficient of the spheres were assigned 3.06 MPa and 2.0 s/m respectively to make the energy return similar to the heel region of the human foot in an athletic shoe [41].

**Table 1:**
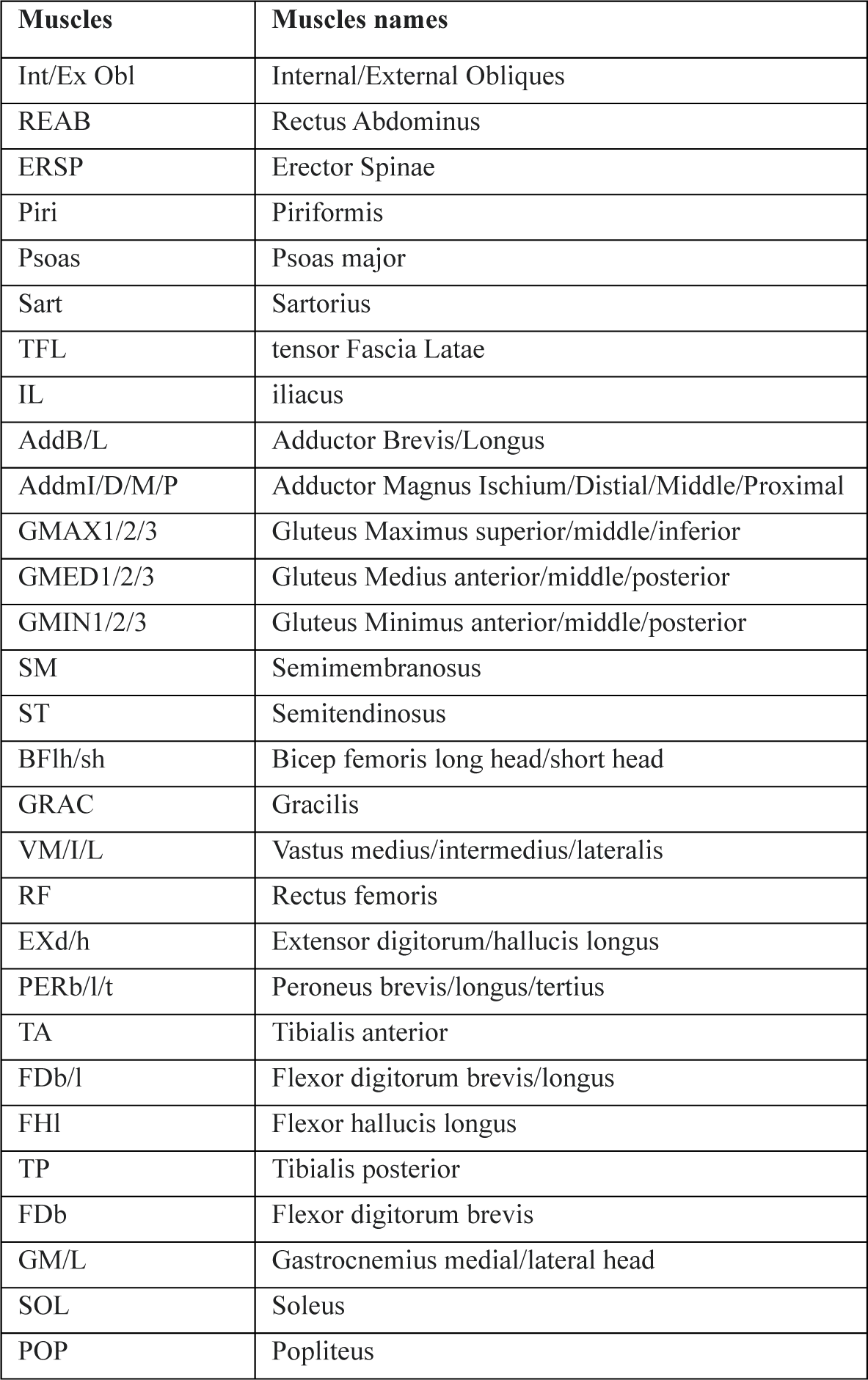
List of the muscles included in the model.

**Table 2:**
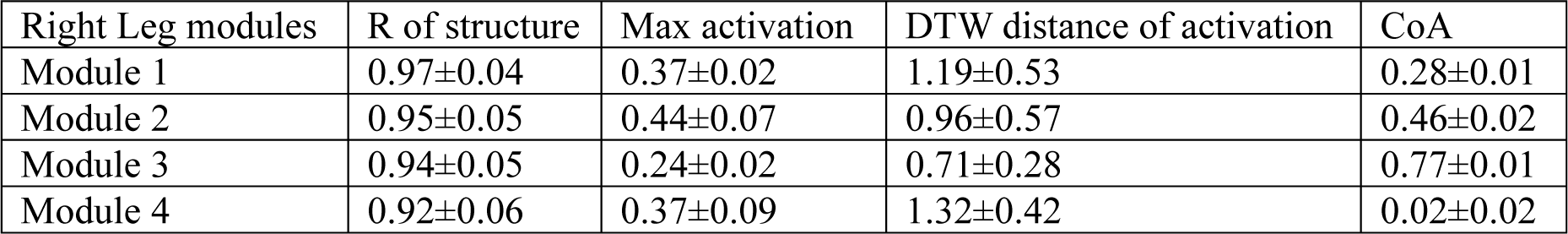
Motor module structure and timing comparison.

### 2.2. Optimal Control Problem and Objective Function

Each of the 27 different desired walking behaviors was generated via a trajectory optimization problem carried out in OpenSim Moco[39]. A 1s stride cycle was discretized into a total of 101 grid points adopting Hermite-Simpson transcription and the discretized problem was turned into a generic nonlinear programming problem using CasADi. This problem took the general form of:

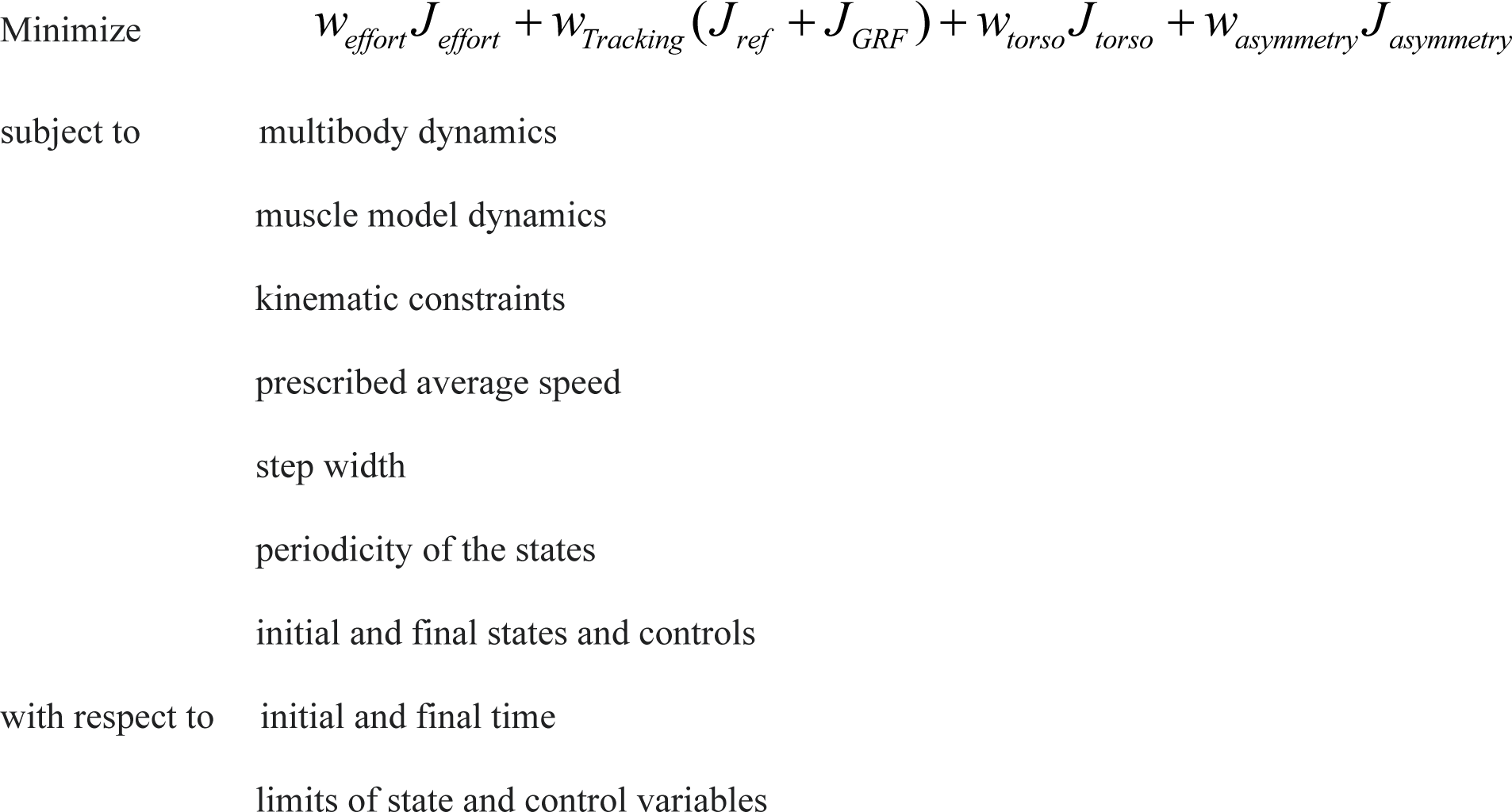

where w’s are the weights of each term to be minimized and each J refers to a cost term that is described in more detail in the below sections. Desired step length asymmetries were achieved via the objective function (J_asymmetrty_) whereas desired speeds and step widths were achieved via constraints.

#### 2.2.1 Step Length Asymmetry goal (J_asymmetry_)

The step length asymmetry term was modeled using the MocoStepLengthAsymmetryGoal and was part of the objective function to be minimized. Step length asymmetry was computed as (RSL-LSL)/(RSL+LSL), where RSL and LSL are right and left step length, respectively. Positive values correspond to larger right step lengths. This goal requires the target asymmetry and total stride length to be prescribed. Simulations of non-asymmetric walking did not include this term and the asymmetry value was set to 0.15 and 0.30 to achieve 15% and 30% step length asymmetry. The total stride length depended on walking speed and was set to 0.74, 1.04 and 1.4 for 0.8, 1.1, and 1.45 m/s, respectively.

#### 2.2.2. Speed constraint

Desired walking speed was achieved via the MocoAverageSpeedGoal. Use of this goal requires a desired value for the average speed of the center of mass and is implemented as a constraint such that the difference between desired and actual walking speed is zero, calculated as:

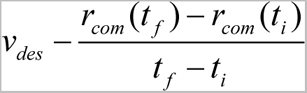

where v_des_ is the desirable speed and the r_com_ is the center of mass position, and t_i_ and t_f_ represent initial time and final time.

#### 2.2.3. Step width constraint

Step width was modeled as a distance constraint between the right and left foot (calcaneus and toe segments) via the MocoFrameDistanceConstraint. This constraint was utilized to keep the minimum sagittal-plane distance between the right and left foot segments greater than a prescribed value. The minimum distance between the feet in normal walking was 0.1m and the step width constraint was used to produce wider steps of 0.2m and 0.3m.

#### 2.2.4 Effort term (Jeffort)

Minimizing effort was modeled as minimizing the sum of the squared control signals (either muscle activation or torque) over the entire simulation via the MocoControlGoal:

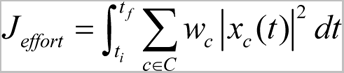

where x_c_ represents each control signal including the 94 MTUs and 3 torque actuators and w_c_ their respective weights, which were set equal to one.

#### 2.2.5. Tracking terms (JRef, JGRF, and Jtorso)

To achieve a realistic walking motion, three different tracking terms were included to track (1) reference kinematics, (2) reference ground reaction forces, and (3) an upright torso orientation.

1. <u>Kinematic tracking</u> was implemented using the MocoStateTrackingGoal, which minimizes the error between simulated joint angles and their reference trajectories (i.e., experimental data). The cost term was defined as [41]: 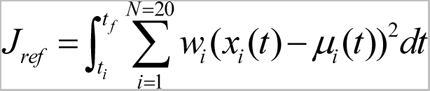 Where x_i_ is the value of i^th^ degree of freedom and µ_i_ is the average experimental value across subjects at time point *t*, and N is the number of DoF being tracked (23 DoF minus 3 trunk angles). The individual tracking weight for each joint was:

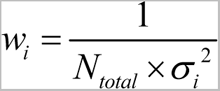 Where σ*_i_* is the standard deviation of experimental data across all subjects of the i^th^ joint averaged over the gait cycle and N_total_=29 (23 plus 3 components of ground reaction forces per leg)
2. <u>Ground reaction force tracking</u> was implemented using the MocoContactTrackingGoal, which minimizes the error between simulated ground reaction forces and their reference trajectories. The cost term was defined as: 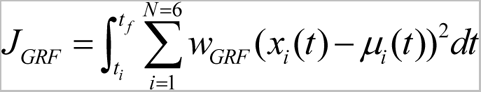 Where *i* refers to i^th^ force component including vertical, anterior-posterior and medial-lateral for both legs. The weight associated with each component was: 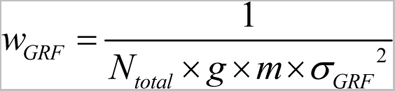 Where g is gravitational acceleration, m mass of the model, σ*_G R F_* is the standard deviation of ground reaction forces across subject from the experimental data at time point *t*.
3. <u>Upright torso orientation</u> was implemented using the MocoOrientationTrackingGoal, which minimized the error between the 3 lumbar rotational DoF and an upright torso orientation. The cost terms was defined as: 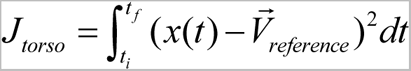 and weight equal to: 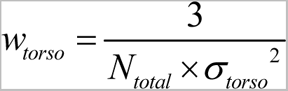 Where *σ_torso_*is standard deviation of between-subjects experimental data of torso orientation.

#### 2.2.5. Periodicity and initial and final state and controls

A periodic gait cycle was produced via the MocoPeriodicityGoal, which imposes equality between initial and final variables for all joint angles and muscle excitation except for pelvis anterior-posterior translation. Furthermore, the initial and final time of the simulation is prescribed such that the whole stride cycle is achieved over a 1 s duration.

### 2.3. Initial Guess Strategy

Twenty-seven distinct walking patterns were created by systematic deviation from experimental data of walking at 1.45m/s from [43]. First, a solution for walking at 1.45 m/s with 0.1m step width and symmetric step length was generated with kinematic tracking from experimental data [43]. This solution served as the initial guess for the optimization of symmetric walking at 1.1m/s, which was then used as an initial guess for the optimization of symmetric walking at 0.8 m/s. Once simulations of symmetric walking with 0.1m step width were achieved at all 3 different speeds, these solutions were used as the kinematic reference data and as the respective initial guess to achieve simulations with 15% asymmetry for each speed, which then served as the initial guess to achieve the 30% asymmetry solution. Then a similar approach was used for generating the 20cm and 30cm step width simulations at each speed and asymmetry level by changing the distant constraint values between feet.

### 2.4 Motor module extraction

Motor modules were identified from each leg for each of the 27 different simulations via nonnegative matrix factorization (NMF)[44]. NMF decomposes muscle activity into a reduced set of motor modules according to the following formula:

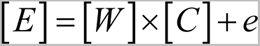

where E is muscle activation matrix of size *m×101* which represents number of muscles on each leg and number of grid points respectively. Matrix W represents motor modules (i.e., the groups of co-active muscles) and is of size *m×n*, where *n* is the number of modules. Matrix C is composed of the activation profiles of each of the *n* motor modules activations and is of the size *n×101*. e is the error of motor module extraction.

### 2.5. Motor module analysis

The following analyses were performed on (1) all muscles per leg and (2) from a reduced experimental subset per leg. This approach enabled us to compare our findings with existing literature, primarily derived from experimental data. The full muscle set included all muscles except the torso muscles due to the inclusion of the torso actuator, resulting in 43 muscles per leg. In contrast, eight muscles per leg were included in the experimental subset: tibialis anterior, soleus, medial gastrocnemius, vastus medialis, rectus femoris, medial hamstrings, lateral hamstrings, and gluteus medius [24,26].

#### 2.5.1 Motor module number

To test our prediction that motor module number does not change with different walking biomechanics, the number of motor modules for each of the 27 different simulated walking behaviors were chosen as follows. For each data matrix (i.e., the *m x 101* E matrix for a single leg within a single simulation), one to 10 (8 for the experimental subset) motor modules (W’s) were extracted. The goodness of fit between actual and reconstructed muscle activity was evaluated with the variability accounted for (VAF), defined as:

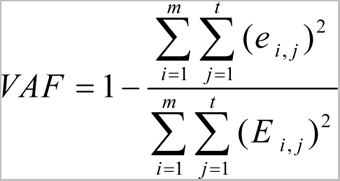

Where *t* and *m* represent number of time points and number of muscles, *E* is muscle activation from the simulations, and *e* represents the error of motor module extraction for each muscle at each time point (EMG reconstructed – *E*).

We chose the number of motor modules, n, for each simulation such that they could account for at least 95% of the patterns of muscle activation, i.e., > 95% VAF. Recognizing that this threshold is an arbitrarily chosen value and there lacks a universally accepted VAF cut-off for choosing motor module number, we further investigated the effect of different thresholds by assessing motor module number with thresholds of 90 and 95%.

#### 2.5.2. Motor module structure

To test our prediction that motor module structure does not change with different biomechanics, two different analyses were performed. For each analysis, the simulated walking of speed 1.1 m/s, step length asymmetry of 0%, and step width of 0.1 m were chosen as the reference solution.

1. The W’s from each solution were compared to those of the reference solution using Pearson’s linear correlation.
2. We also evaluated the ability of the Ws from the reference solution to explain muscle activation in the other simulations. The motor modules activation (Cs) reconstruction were carried out as a non-negative least square problem 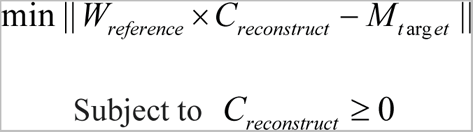 This was then solved using a nonnegative optimizer. Once reconstructed the goodness of the reconstruction was evaluated using VAF. Higher VAF higher the ability of the Ws of the reference solution to explain other walking behaviors. Note that the Cs of each these solutions were compared to assess differences in recruitment timing (Details in Section 4.3).

In addition, the similarity between right and left leg motor module structure due was examined using Pearson’s linear correlation.

#### 2.5.3. Motor module recruitment timing

To explore how motor module recruitment varied across different walking behaviors, the reconstructed Cs (as described above in 2.5.2) were compared across simulations using three different outcome metrics:

1. <u>Maximum activation</u> to assess differences in peak module recruitment.
2. <u>Center of activity</u> (CoA) to assess differences in recruitment timing. CoA of each motor module activation was computed using circular statistics[45]. 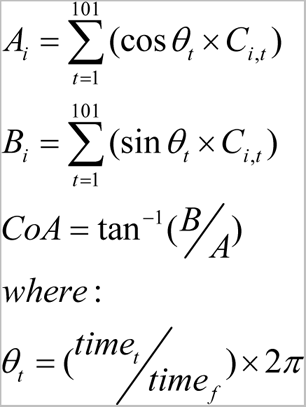 Here, *i* represents the *i*^th^ motor module activation for which the CoA calculation is carried out and *t* the time points. θ_t_ is the phase of the gait cycle which was derived by multiplying the normalized time vector and two radians accounting for the whole stride cycle.
3. <u>Dynamic Time Warping</u> (DTW) to assess similarity of each recruitment timing curve to that from the reference solution. DTW is a well-known algorithm to find the optimal alignment between timeseries and has found success in comparing biomechanical timeseries during gait such as ground reaction forces, muscle activations and joint angles[46–48]. DTW assesses similarities between two timeseries after aligning for temporal distortions, producing a measurement of similarity between motor module recruitment timing curves irrespective of overall timing differences (e.g., heel-strike occurring at different timepoints) and is thus a good complement to CoA. DTW analysis in this study was carried out using tha DTW built-in MATLAB function. Lower values of DTW are associated with higher similarity.

## 3. Results

### 3.1. Target Gaits

All 27 simulations converged to a stable solution for a 1 s stride cycle. Constraints for step width (0.1, 0.2 and 0.3m; Fig. 1A) and average walking speed (0.8, 1.1 and 1.45 m/s; Fig. 1B) were achieved and target asymmetry levels (0, 15 and 30%) were within 1% accuracy.

**Figure 1:**
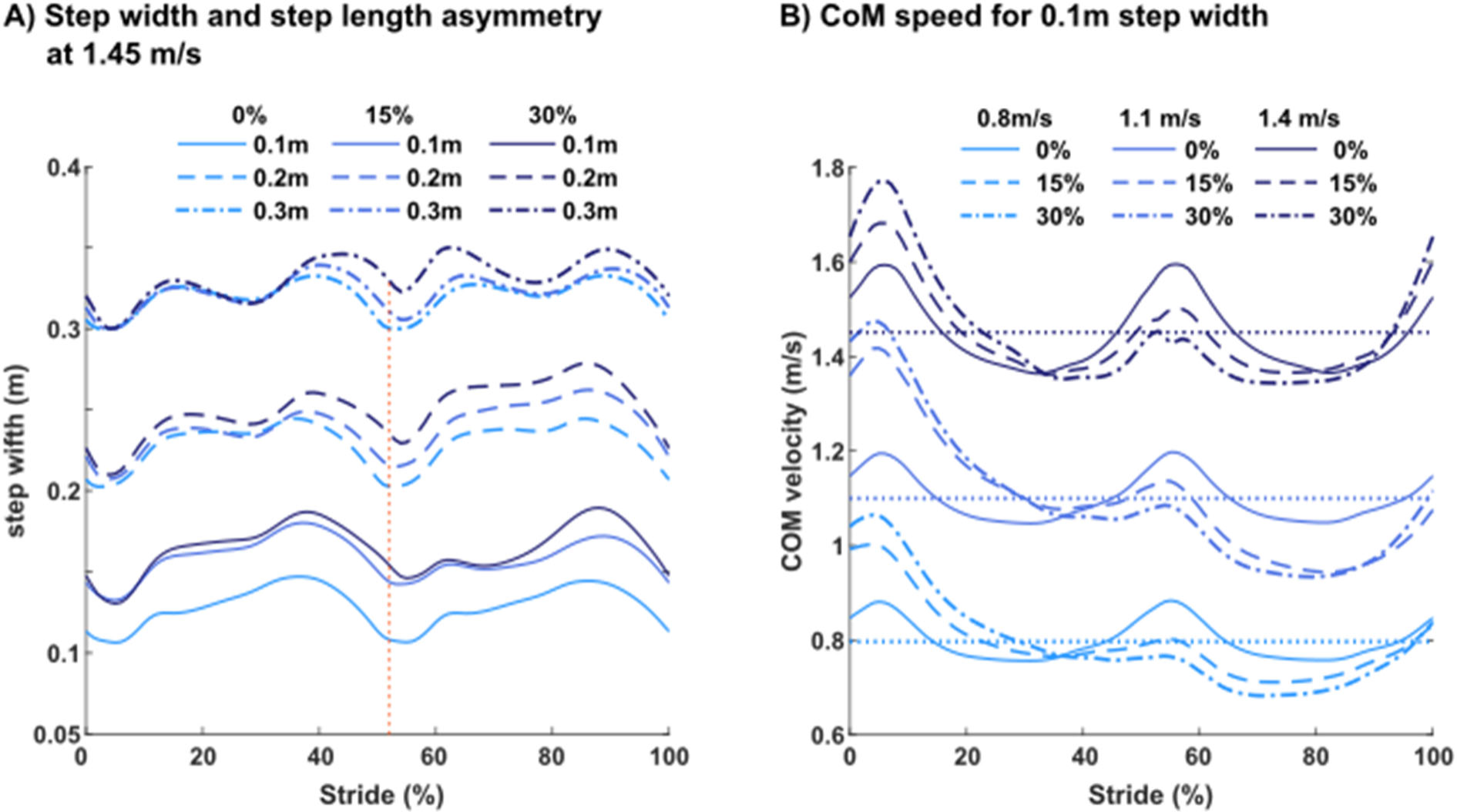
A) Simulations of walking at 1.45 m/s with different step length asymmetry levels demonstrating the different step widths achieved. The minimum step width occurred at heel-strike (red dotted line) B) Simulations of 0.1m step width with different asymmetry levels demonstrating center-of-mass (CoM) speed over the gait cycle and it’s average (i.e., the target value for walking speed).

### 3.2. Motor Module Number

Motor module numbers were similar across simulations. When extracting motor modules from the experimental 8-muscle set with a 95% VAF cutoff, the number of motor modules was 3 for all walking behaviors in both legs (Fig. 2). Motor module number was increased and slightly more variable when extracting from the full muscle set. The median number of motor modules to achieve the 95% VAF threshold was 5 for both legs but ranged between 4-6 modules (Fig. 3B; left leg: range 4-5, 4.33±0.47; right leg: range 4-6, 4.81±0.55). There was a trend for an increase in motor module number with more complex gait biomechanics (i.e., faster walking speeds + more asymmetry + wider steps). However, the difference between whether 4 or 5 modules was needed came down to differences in VAF of 3% or less (Fig. 3A). When using the 90% VAF threshold to choose motor module number for the full muscle set, motor module numbers were more consistent across simulations with the median number of motor modules equal to 3 for both the left and right legs (Fig. 3C; left leg: range 3, 3±0; right leg: range 3-4, 3.15±0.35).

**Figure 2.**
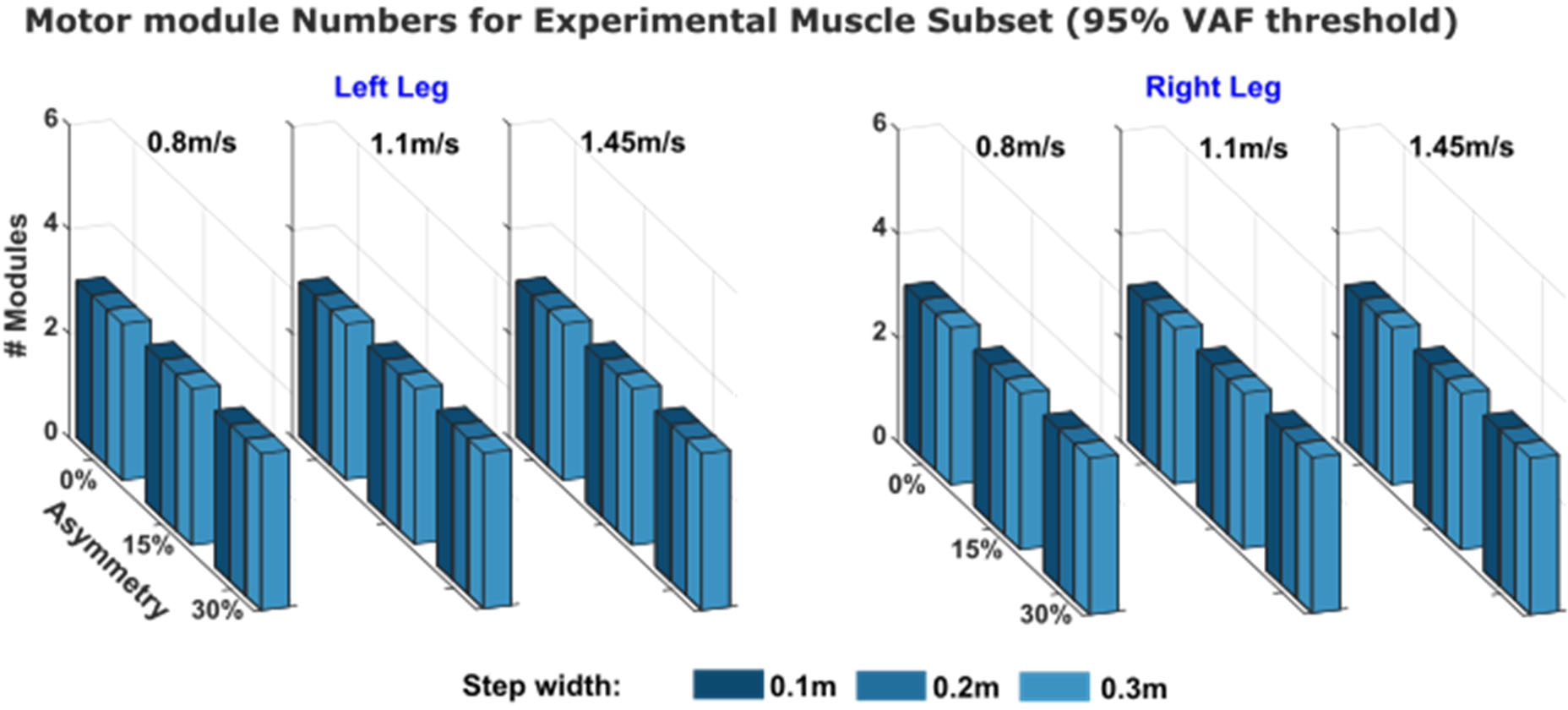
Number of motor modules to achieve 95% VAF for each simulation using only the 8-muscle experimental subset.

**Figure 3.**
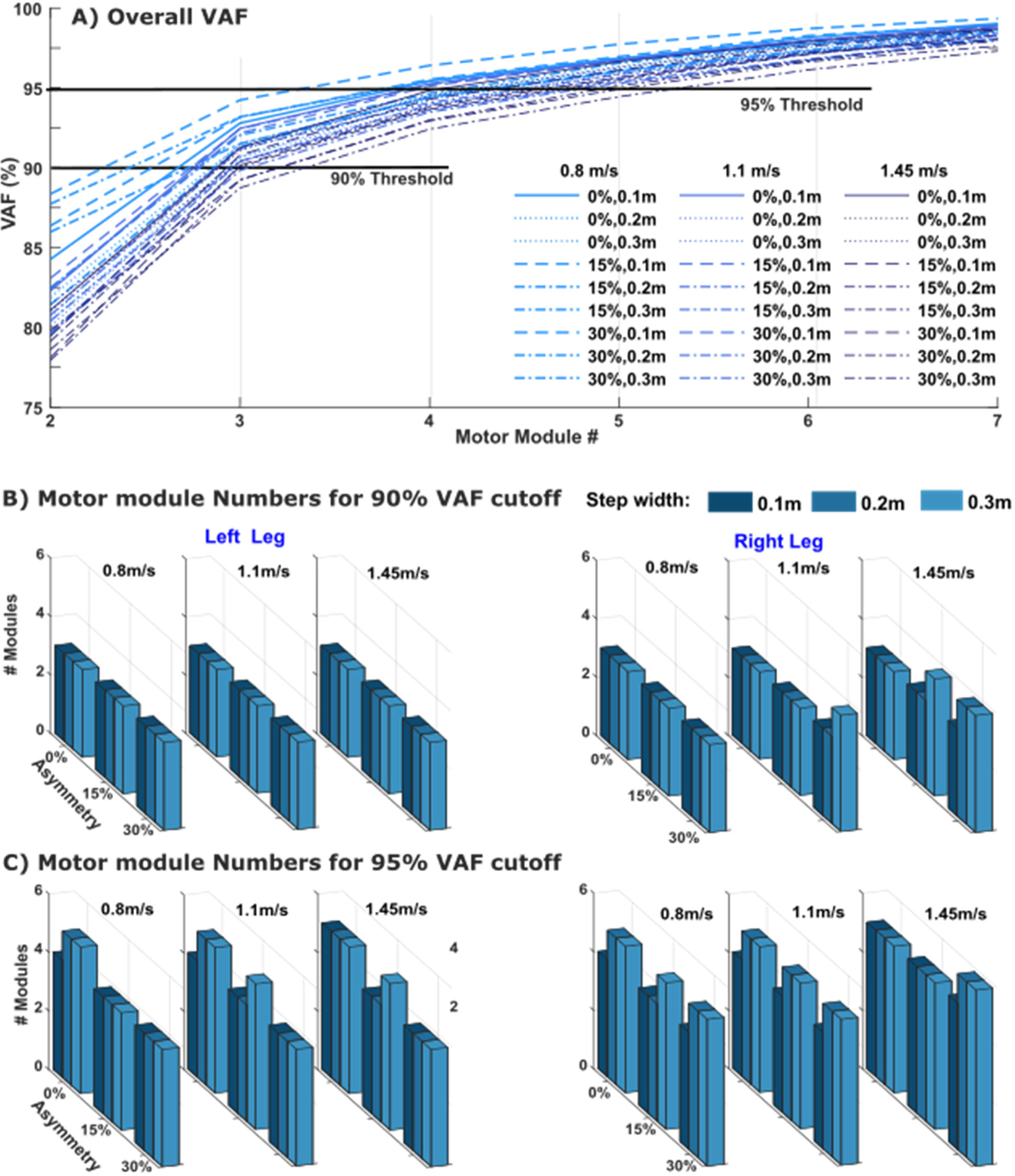
A) Variability accounted for (VAF) in each simulation for each number of motor modules for all muscles on the right leg. B) The number of motor modules required for each simulation to reach the 90% VAF threshold. C) The number of motor modules required for each simulation to reach the 95% threshold.

### 3.3. Motor Module Structure

Motor modules with similar structure were recruited across the simulations with different biomechanics. Fig. 4A shows a histogram of the similarity in motor module structure across all simulations in both legs. The correlation coefficient of the motor modules from each simulation compared to those from the reference solution (i.e., symmetric walking at 1.1 m/s with 0.1m step width) extracted from the full muscle set was 0.94±0.05 and 0.91±0.07 for the right and the left leg, respectively. When extracting motor modules from the experimental muscle subset, the similarity in module structure was 0.99±0.02 and 0.98±0.02 for the right and the left leg, respectively.

**Figure 4:**
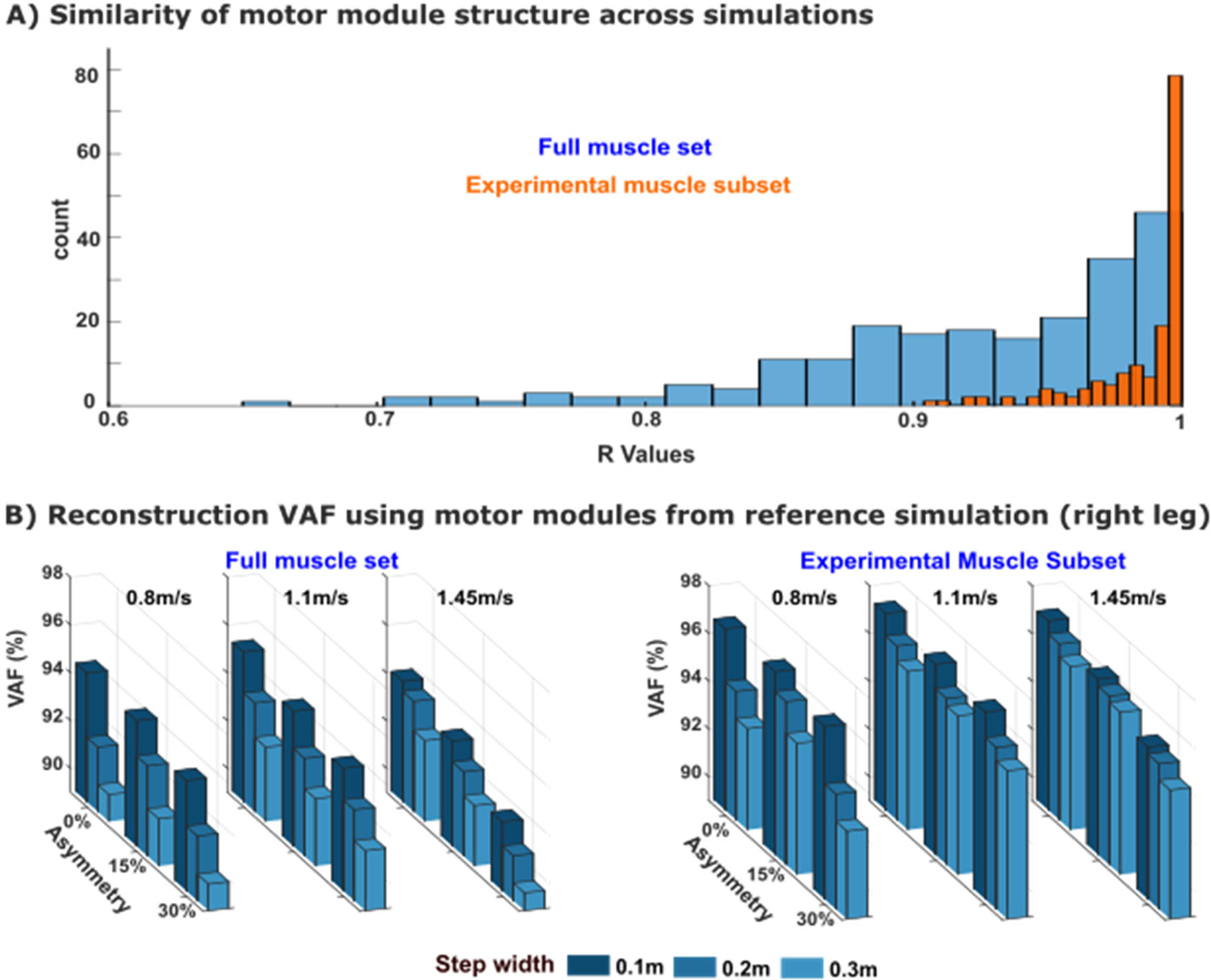
A) Similarity in motor module structure using the motor modules extracted from each individual simulation for both the experimental muscle subset (orange) and full muscle subset (blue) for both legs. B) The variability accounted for (VAF) of muscle activity in the right leg when using motor modules from the reference simulation to reconstruct each of the other simulations.

The similarity of motor module structure across simulations with different walking biomechanics was confirmed when using the motor modules from the reference solution to reconstruct muscle activity in each of the other simulations (Fig. 4B). Reconstructed motor modules explained muscle activity of all simulations with at least 90% VAF (0.93 ±0.01 for either leg). The lowest VAF reconstruction of 90% was walking at 1.45m/s with 0.3m step width and 30% step length asymmetry, i.e., one of the most complex walking behaviors simulated.

Motor modules for the right leg of the reference simulation (1.1 m/s, 0.1m step width, and 0% step length asymmetry) are illustrated in Fig. 5. Module R1 consisted of the glutei, knee extensors, and some hip flexors. This module was activated throughout much of the stance phase (Fig. 7-9). Module R2 consisted of the plantar flexors (gastrocnemius and soleus) with additional representation from the iliacus and psoas. This module was active during late stance. Module R3 consisted primarily of hip flexor muscles with additional activity from the hamstrings and ankle dorsiflexors. This module was active from late stance into swing. Module R4 consisted primarily of the ankle dorsiflexors and was active in late swing into early stance. The motor modules for the experimental muscle subset (Fig. 5B) were similar to those extracted from the full muscle set, with the exception that Module R3 was not identified due to, perhaps, the lack of hip flexor muscles in the experimental dataset.

**Figure 5:**
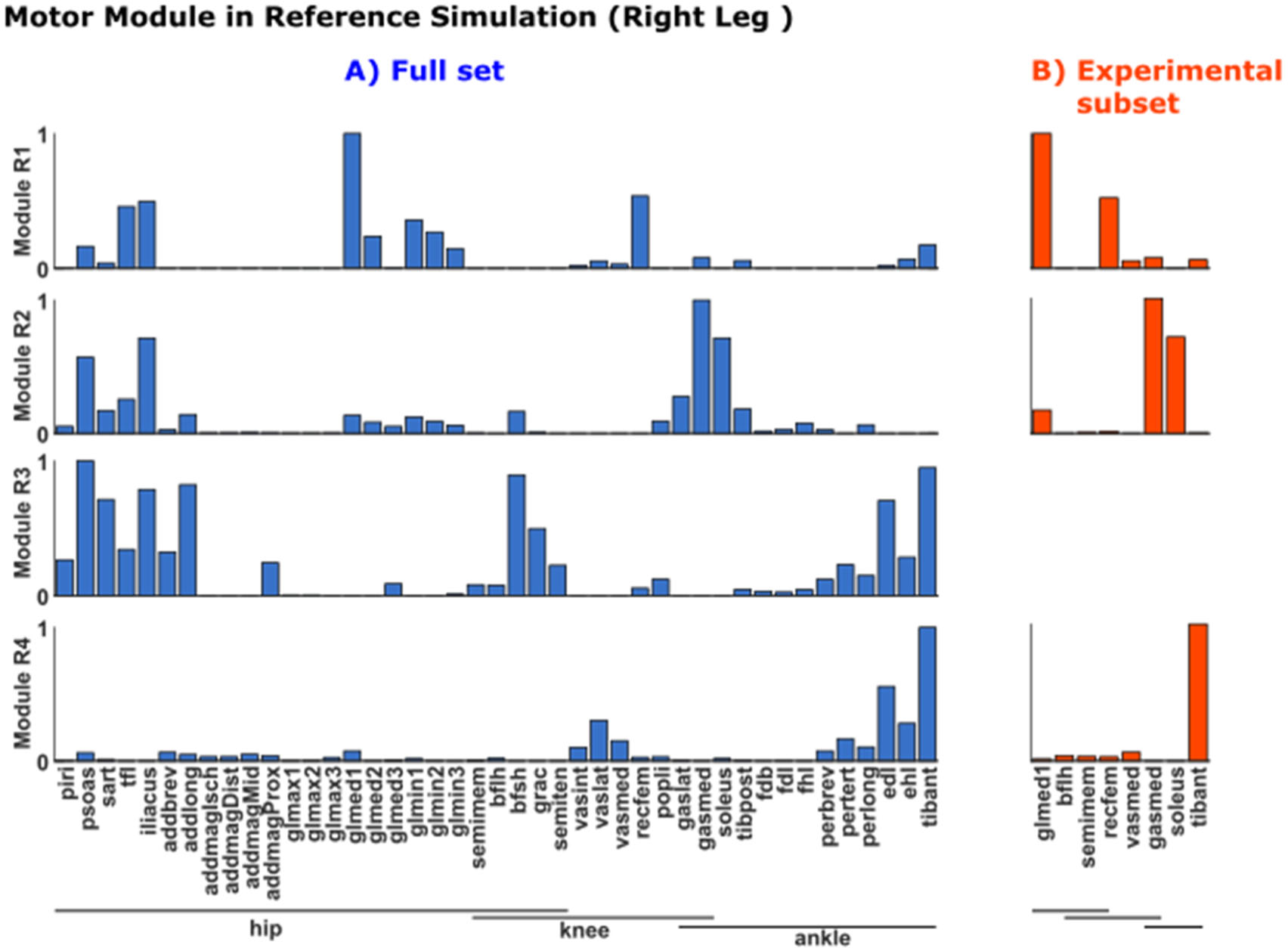
Motor modules on the right leg in the reference simulation (1.1 m/s, 0.1m step width, and 0% step length asymmetry) for (A) the full muscle set and (B) the experimental muscle set.

Motor module structure also did not substantially differ between legs. As expected, there were slight differences due to step length asymmetry (Fig. 6). These asymmetry-related differences increased with speed and occurred alongside larger differences between absolute step lengths in each leg that also increased with speed. Nevertheless, most modules remained over 80% similar between legs.

**Figure 6:**
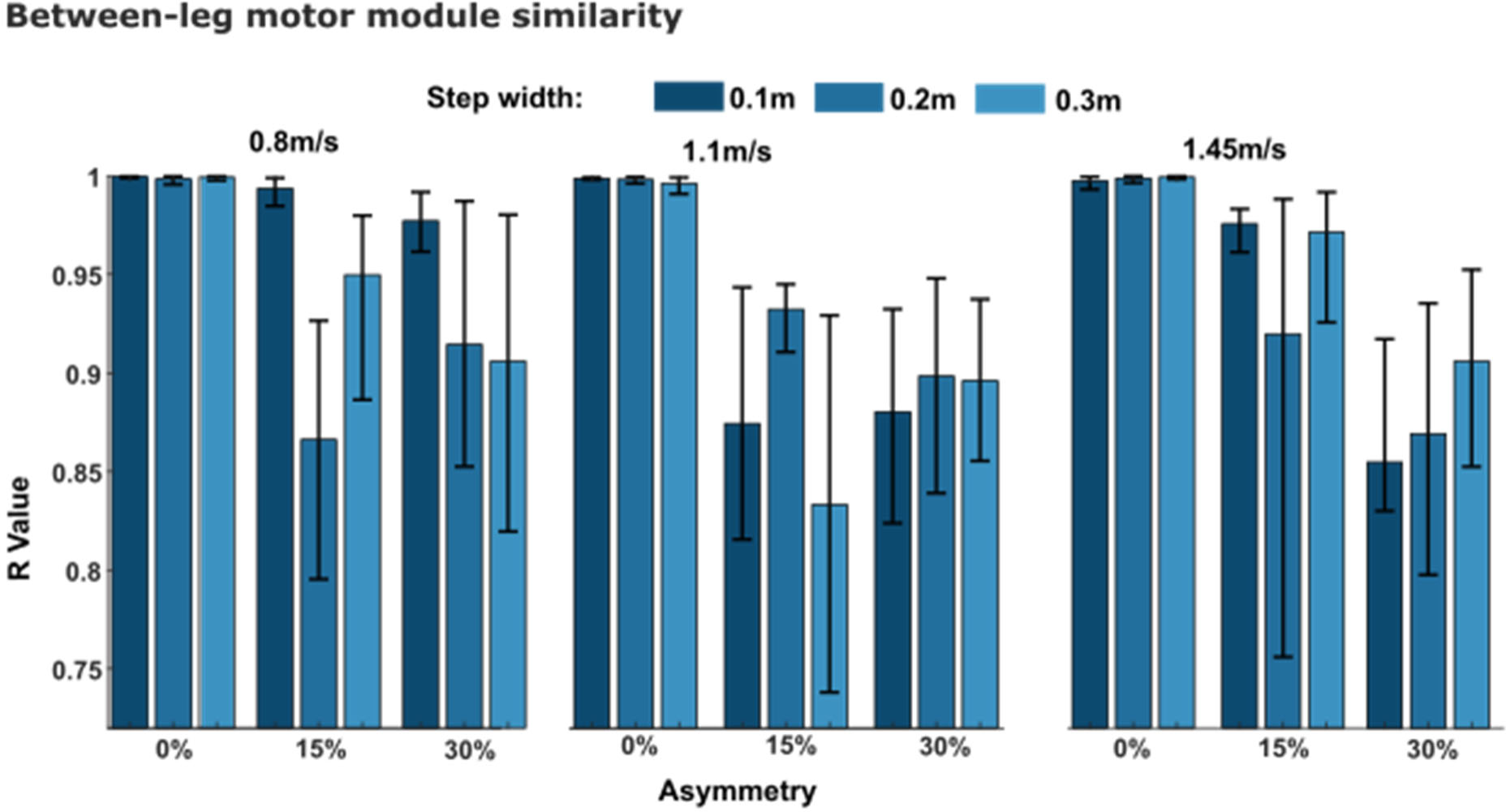
Similarity in motor module structure between legs in each simulation. The bars represent the average value of module comparisons and the error bars represent the minimum and maximum value of similarity.

### 3.4 Motor Module Recruitment Timing

Here, we compared the motor module activation patterns (i.e., the C’s) when using the reference solution motor modules (i.e., the W’s) to reconstruct muscle activity for each simulation. Peak activation and DTW values differed across solutions (Table 1, Figs. 7-9), but CoA remained largely unchanged. Specific differences in recruitment timing curves as a function of walking speed, step length asymmetry, and step width are presented in the below sections.

**Figure 7:**
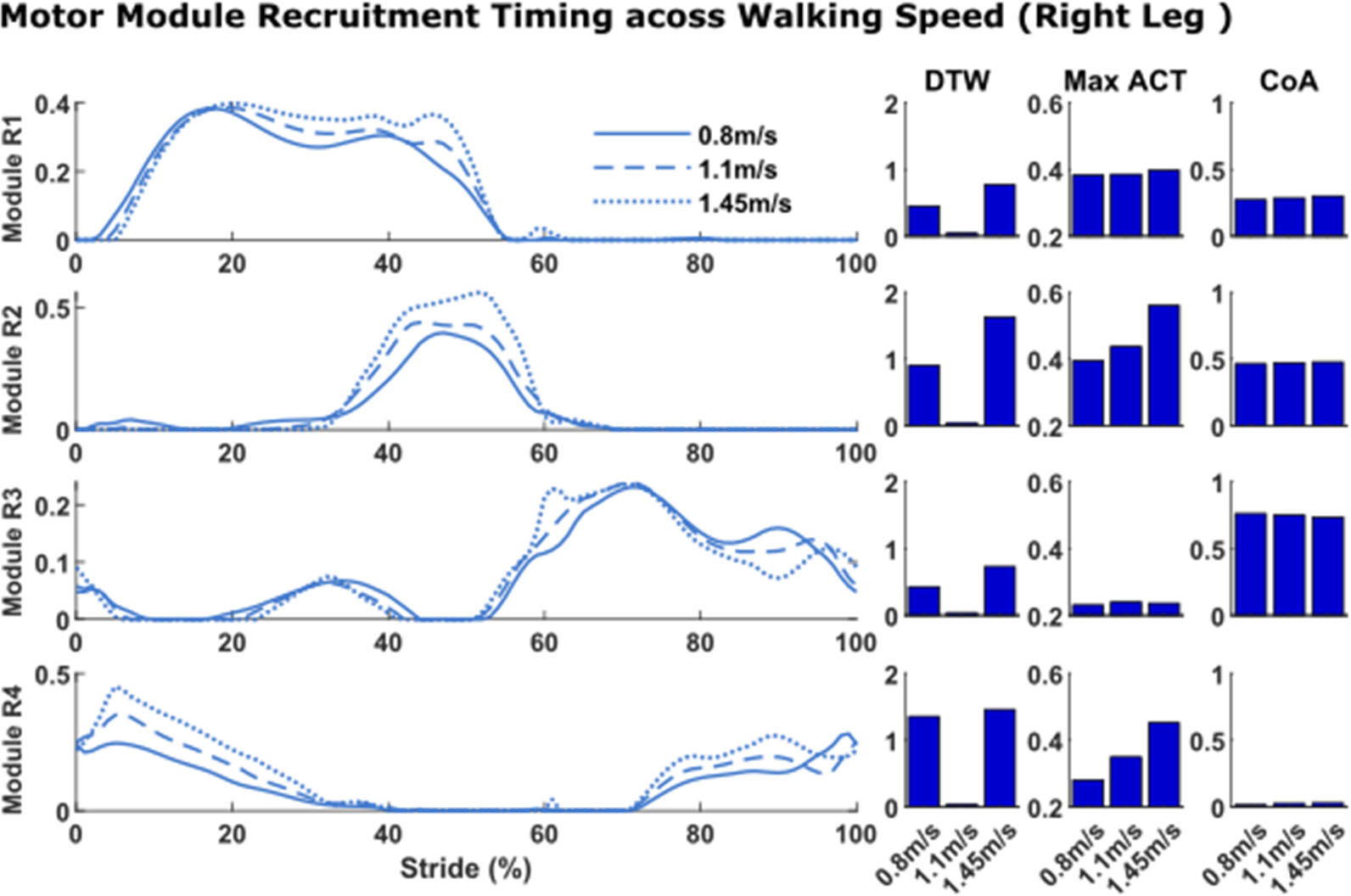
Changes in motor module recruitment timing that occurred with walking speed. The solutions illustrated are those with 0% step length asymmetry and 0.1m step width.

**Figure 8:**
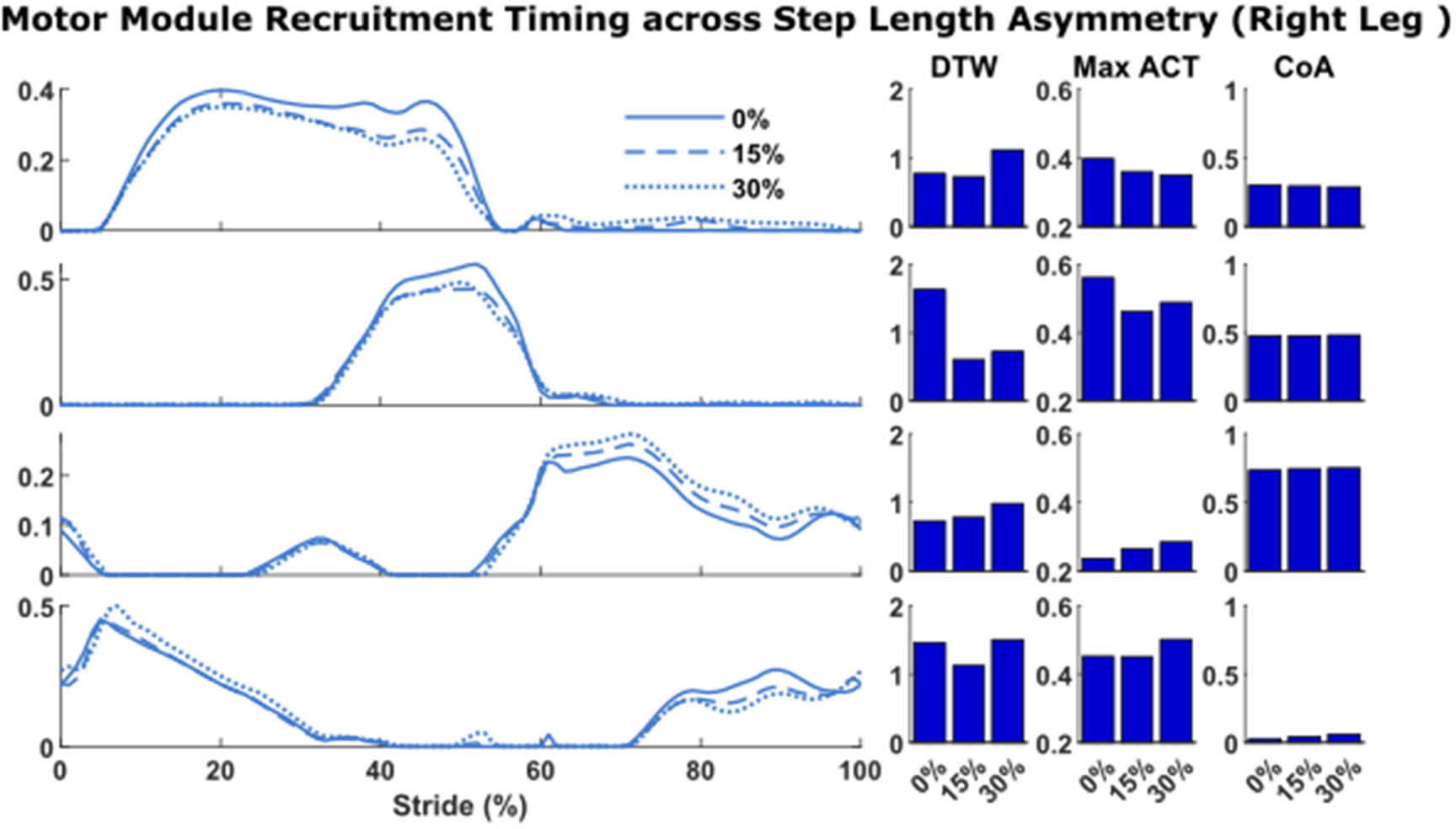
Changes in motor module recruitment timing that occurred with step length asymmetry. The solutions illustrated are those with walking speeds of 1.45m/s and step width of 0.1m.

**Figure 9:**
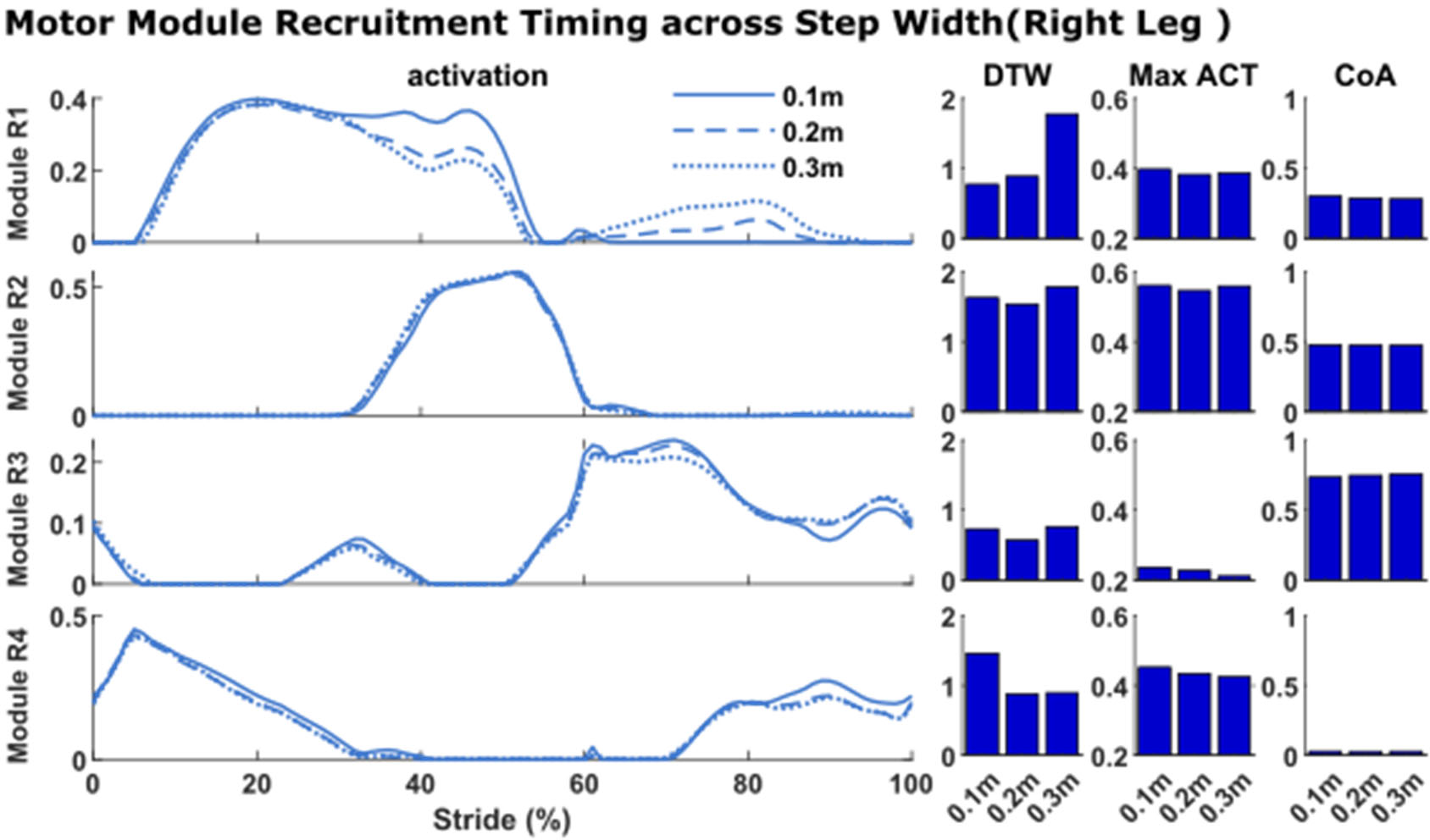
Changes in motor module recruitment timing that occurred with step width. The solutions illustrated are those with walking speed of 1.45m/s and symmetric step lengths.

#### 3.4.1. Effect of walking speed

Motor module recruitment timing changed with walking speed (Fig. 7; solutions for 0.1m step width and 0% asymmetry across speeds). Module R1, which consisted of the glutei and knee extensors, exhibited changes in the pattern of its recruitment timing over the gait cycle as revealed by DTW. Qualitatively, these changes occurred due to peak activity levels extending longer in the stance phase. Module R2, which consisted of the plantarflexors, exhibited both increases in peak activation and changes in recruitment pattern with walking speed. Qualitatively, the peak activity was reached earlier and maintained for longer when walking at faster speeds. Module R3, which consisted primarily of hip flexor muscles with additional activity from the hamstrings and ankle dorsiflexors, exhibited only small differences in timing patterns. Module R4, which consisted primarily of the ankle dorsiflexors, exhibited both increases in peak activation and changes in recruitment pattern with walking speed. Qualitatively, the peak activity was reached earlier in the swing phase and maintained for longer in the stance phase when walking at faster speeds.

#### 3.4.2. Effect of step length asymmetry

Motor module recruitment timing also changed with step length asymmetry (Fig. 8), although to a lesser extent than with speed. The largest changes in recruitment timing across different step length asymmetries occurred when walking at the fastest speed. Peak muscle activity increased in module R3 (hip flexors) and maintained heightened levels throughout swing, whereas modules R1 (glutei and knee extensors) and R2 (plantarflexors) exhibited decreased activity in late stance.

#### 3.4.3. Effect of step width

Motor module recruitment timing exhibited fewer changes across step width solutions (Fig. 9). The largest changes in recruitment timing across different step widths occurred when walking at the fastest speed. Module R1 (which includes both hip flexor and abductor muscles) exhibited changes in timing pattern such that its activation was decreased in late stance and increased throughout swing phase. Recruitment of modules R2-R4 remained largely identical across step widths.

## 4. Discussion

The purpose of this study was to explore the extent to which motor modules are emergent from pathological walking biomechanics versus representative of an underlying neural control strategy or in other words, do pathological walking biomechanics require a reduction in motor module number? Given the complexity of identifying motor modules compared to simply recording movement biomechanics, answering this question is critical to support the increasing use of using motor modules to identify neuromuscular deficits limiting walking function, guide rehabilitation efforts, and control exoskeleton and prosthetic devices. To address this question, we utilized musculoskeletal modeling and simulation to evaluate the influence of various characteristics of pathological walking biomechanics on the number and structure of motor modules that explain muscle activity in the lower limbs. This approach allowed us to disentangle biomechanics from neural control, something that is difficult to do experimentally. We found that different walking speeds, step widths, and step length asymmetries could be achieved via the same motor modules. This supports our hypothesis that the differences in motor modules observed between individuals with and without pathological walking function reflect, to some extent, differences in underlying neural control strategies and not just differences in their walking biomechanics.

Motor module numbers were largely unaffected by different walking biomechanics. Three motor modules were required to explain simulated activity of the experimental subset of 8 muscles per leg in every simulation (Fig. 2). This number is consistent with prior studies that have identified between 3-5 motor modules in healthy adults [24,49–51]. Motor module numbers in the current study are likely on the lower end because (1) each simulation consisted of only a single gait cycle and (2) simulated muscle activity is less contaminated by noise than experimental EMG, both of which are known to lead to fewer motor modules [22,52–56]. As expected and consistent with prior studies [57,58], motor module numbers increased when extracting from the full set of 43 muscles per leg. Moreover, the consistency of motor module numbers across simulations was highly dependent on the criteria used to select motor module numbers. Motor modules were identical across simulations when using VAF cutoff of 90% but became slightly more variable across simulations when using VAF cutoff of 95%. Selecting the appropriate criteria for identifying motor module numbers has historically been a challenge, and it is well-recognized that these somewhat arbitrary cutoffs can impact results [22]. The increased variability in motor module numbers observed with a stricter selection criterion likely stemmed from reconstructing activity in muscles that play a minor role in locomotion, as muscle activity tends to be more variable when not substantially contributing to task demands [59–62]. As a secondary analysis of motor module numbers, we assessed how well the motor modules from the reference solution could reconstruct muscle activity from the other simulations. We found that over 90% VAF could be explained in all simulations (Fig. 4B), confirming that the same number of motor modules could reconstruct muscle activity across simulations across simulations with various characteristics of pathological walking biomechanics contrasts. These results contrast with previous experimental studies in which motor module number was reduced in those with pathological function (e.g., cerebral palsy, Parkinson’s disease, stroke, etc.) [7,24,27–32] and provide evidence that such a reduction in motor module number is not solely attributable to altered biomechanics but also likely involves a neural component (e.g., neural injury, altered neural control strategy, etc.).

Motor module structure was also largely unaffected by different walking biomechanics. The motor modules extracted from the full muscle set in the reference simulation (i.e., “normal” walking biomechanics) (Fig. 5) were similar to those identified in other studies in which there were modules with dominant activity from: (1) hip/knee extensors in early stance, (2) ankle plantar flexors in late stance, (3) hip flexors and hamstrings from late stance into swing, and (4) ankle dorsiflexors in late swing into early stance [24–26,63,64]. Likely due to the absence of hip flexor musculature, motor module 3 was absent from the experimental subset. We found that these same motor modules were present in all simulations regardless of walking biomechanics for both the full muscle set and the experimental subset. The similarity of motor module structure across simulations was greater than 0.9 for all comparisons of motor modules from the experimental subset. Although the similarity was slightly reduced in the full muscle set, most comparisons remained above 0.85 similar, suggesting consistency in motor module structure across simulations. Moreover, that muscle activity could be reconstructed using the motor modules from the reference simulation (Fig. 5) with over 90% VAF in all simulations (Fig. 4B) confirmed the consistency of motor module structure across simulations. The different abnormal walking biomechanics were instead achieved via modulation of motor module recruitment timing. How the timing of each motor module changed across conditions was consistent with their previously identified biomechanical roles [34,35,65]. For example, consistent with their respective role in body support and forward propulsion, modules R1 (hip/knee extensors) and R2 (plantarflexors) increased their activity in early and late stance as walking speed increased (Fig. 7). Consistent with their roles in controlling leg swing, module R3 (hip flexors) exhibited heightened activity with increasing step length asymmetry whereas module R1 (which also included hip adductor musculature) exhibited heightened activity with increasing step widths. Taken together, these results demonstrate that the same motor modules can be used to produce pathological-like walking biomechanics, providing evidence that alterations in motor module structure that occur in pathological populations have at least some neural origins.

While our study provides valuable insights into the role of biomechanics versus neural control in motor module structure and recruitment during walking, there are several limitations to consider. First, we investigated only a limited number of spatiotemporal characteristics of walking biomechanics: walking speed, step width, and step length asymmetry. This choice was motivated by the fact that these spatiotemporal characteristics are commonly altered in individuals with neurological disease/injury and accompanied by a reduction in motor module number [7,24,27–32]. Second, our cost function included a term to track experimental kinematics from healthy walking to generate our initial simulations at each speed. However, the value of the weight assigned to this term allowed the solution still to deviate substantially from the tracked kinematics to achieve all desired abnormal spatiotemporal targets.

Moreover, including the tracking term also helped to prevent unwanted changes in walking biomechanics other than the desired changes in speed, step length asymmetry, and step width. Lastly, we used generic musculotendon parameters (e.g., muscle strength, tendon slack length, etc.) even though these parameters may change in individuals with neurological disease or injury. However, prior simulation studies have shown that impaired musculotendon properties alone do not prevent normal walking biomechanics and that pathological walking biomechanics only emerged when combined with reduced motor module number and/or the goal to minimize energy expenditure [37,66]. Therefore, we do not expect our main results demonstrating that pathological walking biomechanics can be achieved without a reduction in motor module number to be affected by this limitation. Future work should further explore the interaction between motor module number, musculotendon parameters, and optimality to identify what combinations of neural constraints and neural strategies might lead to pathological walking biomechanics.

## 5. Conclusions

This study used predictive simulations to explore the role of pathological walking biomechanics on the number and structure of motor modules to explain lower limb muscle activity. We identified a consistent set of motor modules across simulations to muscle activity across varying walking speeds, step widths, and step length asymmetries. These results differ from experimental studies in which these same spatiotemporal walking biomechanics are associated with alterations in EMG-derived motor modules.

Consequently, our results emphasize the significant impact of neural deficits and neural control strategies on alterations in EMG-derived motor modules in pathological populations. These insights provide support for the potential use of EMG-derived motor modules in identifying neuromuscular deficits, guiding rehabilitation interventions, and controlling exoskeletons and prosthetic devices in individuals with pathological walking function.

## Declarations

### Ethics approval and consent to participate

Not applicable

### Consent for publication

Not applicable

### Availability of data and materials

All results and the codes necessary to replicate the simulations and extraction of motor modules are available at: https://github.com/Mohammad-Rahimi/different_walking_behaviors_directCollocation_NMF

### Authors’ contributions

JLA: conceptualization, study design, data interpretation, writing – reviewing & editing, funding acquisition, project administration. MRG: study design, formal analysis, data interpretation, writing – original draft, visualization, writing – reviewing & editing. All authors contributed to the article and approved the submitted version.

## List of abbreviations

CoA: center of activity
DTW: dynamic time warping
DoF: degree of freedom
EMG: electromyography
MTU: muscle-tendon unit
NMF: nonnegative matrix factorization
VAF: variability accounted for

## Acknowledgements.

Not applicable.

